# Quantitative fluorescence emission anisotropy microscopy for implementing homo-FRET measurements in living cells

**DOI:** 10.1101/2022.10.01.510443

**Authors:** Thomas S van Zanten, Greeshma Pradeep S, Satyajit Mayor

## Abstract

Quantitative fluorescence emission anisotropy microscopy reveals the organization of fluorescently labelled cellular components and allows for their characterization in terms of changes in either rotational diffusion or homo-Förster’s energy transfer characteristics in living cells. These properties provide insights into molecular organization, such as orientation, confinement and oligomerization *in situ*. Here we elucidate how quantitative measurements of anisotropy using multiple microscope systems may be made, by bringing out the main parameters that influence the quantification of fluorescence emission anisotropy. We focus on a variety of parameters that contribute to errors associated with the measurement of emission anisotropy in a microscope. These include the requirement for adequate photon counts for the necessary discrimination of anisotropy values, the influence of extinction coefficients of the illumination source, the detector system, the role of numerical aperture and excitation wavelength. All these parameters also affect the ability to capture the dynamic range of emission anisotropy necessary for quantifying its reduction due to homo-FRET and other processes. Finally, we provide easily implementable tests to assess whether homo-FRET is a cause for the observed emission depolarization.

## Introduction

Fundamental processes of intracellular life may be understood in terms of pools of molecules that diffuse, interact, bind, change conformation and catalyse physical and chemical state changes within the confined volume of a cell. Utilizing the arsenal of available techniques allows the researcher to unravel rules that govern these processes (Zanten and Mayor, 2015). Traditional biochemical methods (grind and find) provide ensemble averages of concentrations as well as information on interacting species, but lack access to spatial information at the scale of the processes carried out inside the cell. Chemical specific spatial contrast is achieved by using tagged markers in combination with imaging. In a fluorescence setting, optically resolved spatial information can be coupled with high temporal resolution and high signal-to-noise in living systems. However, far-field based fluorescence techniques are inherently diffraction limited and following individual proteins at a density above 10 per um^2^ becomes difficult (Jaqaman *et al*., 2008; Manley *et al*., 2008; Sergé *et al*., 2008). Therefore, it remains challenging to provide direct *in situ* evidence of interaction, binding and conformational changes of molecules in physiological conditions.

Fluorescence resonance energy transfer (FRET) is uniquely sensitive to the detection of short length scale interactions between fluorophores. This allows a window onto events that occur at 1-10 nm range even though one uses diffraction limited imaging systems (Krishnan *et al*., 2001). Depending on several important conditions (such as spectral overlap, the distance between, and the orientation of the fluorophore dipoles) that have been documented previously (Jares-Erijman and Jovin, 2003), two spectrally distinct fluorophores can transfer the energy of the excited state donor (blue-shifted) fluorophore to the acceptor (red-shifted) fluorophore (Hoppe *et al*., 2002) in a non-radiative process called hetero-FRET. This concomitantly occurs with a decrease in the donor fluorophore lifetime (Wallrabe and Periasamy, 2005). The sensitivity of this process to the fluorophore distance at the nanometer scale invokes the adage of a spectroscopic ruler (Stryer, 1978). An increase in FRET is additionally associated with a depolarization of the emission if donor fluorophores are excited with polarized light (Förster, 1948). Especially when donor and acceptor are on different molecules (Rizzo *et al*., 2006) the polarization-based method benefits from a large dynamic range and reduced background from the acceptor fluorophore (Sharma *et al*., 2004; Rizzo and Piston, 2005). However, the ratio between donor and acceptor fluorophores influences the dynamic range and sensitivity of the hetero-FRET process.

FRET between like-fluorophores called homo-FRET also takes place. Instead of measuring the changes in lifetime of the donor or spectral shift of the detected fluorescence, determining the extent of homo-FRET is obtained by measuring the reduction in emission polarisation anisotropy (Weber, 1954; Lidke *et al*., 2003). The homo-FRET methodology eliminates the requirement for careful tuning of donor-acceptor ratios. Consequently, the method is advantageous for detecting homo-oligomerization even when only a very low fraction of interacting species is present (Varma and Mayor, 1998; Sharma *et al*., 2004). This permits a measurement of oligomerization below the detection limit of hetero-FRET labelling techniques (Kenworthy and Edidin, 1998; Sharma *et al*., 2004). Here, we provide a systematic way to perform steady-state homo-FRET measurements, and a comprehensive and practical compendium for the use of emission anisotropy for the detection of homo-FRET. We document factors that will affect the dynamic range of the measurement, measurement-associated errors and suggest ways to attribute the anisotropy measurement to homo-FRET. This will allow a rigorous and quantitative implementation of this powerful technique in biological systems where such resolution is necessary.

### Measuring homo-FRET: Theoretical background

The magnitude of homo-FRET may be determined by measuring the extent of depolarization of fluorescence emission (or reduction in emission anisotropy) upon exciting fluorophores with polarized light (Lakowicz, 2006). In this photophysical process, fluorophore dipoles that are aligned parallel to the plane of polarization of the excitation beam are excited. Fluorescence emission anisotropy, *r*, is detected by collecting emitted photons at two orthogonal polarizations. The following expression is used to define fluorescence anisotropy:

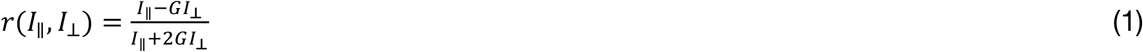

where *I*_‖_and *I*_⊥_ are the detected emission intensities parallel and perpendicular to the excitation, respectively. *G* is the g-factor of the microscope defined as the detection efficiency of the parallel polarized photons with respect to the perpendicular polarized photons. The denominator in equation 1 is equivalent to the total number of photons detected, making it possible to compare anisotropy values from heterogeneous regions. In other words, the detected anisotropy is the fractional sum of anisotropies originating from the region of measurement. For an ensemble of immobile but isotropically oriented fluorophores that have their emission dipole aligned with their absorption dipole (Figure 1Ai) the maximum attainable anisotropy for single-photon excitation is 0.4. Angular mismatch between the absorption and emission dipole, *β*, results in an instantaneous depolarisation and determines the fluorophore dependent intrinsic anisotropy, *r*_0_, that is lower than 0.4 (e.g. GFP; *β* = 11º, *r*_0_ = 0.389 (Myšková *et al*., 2020)).

**Figure 1.**
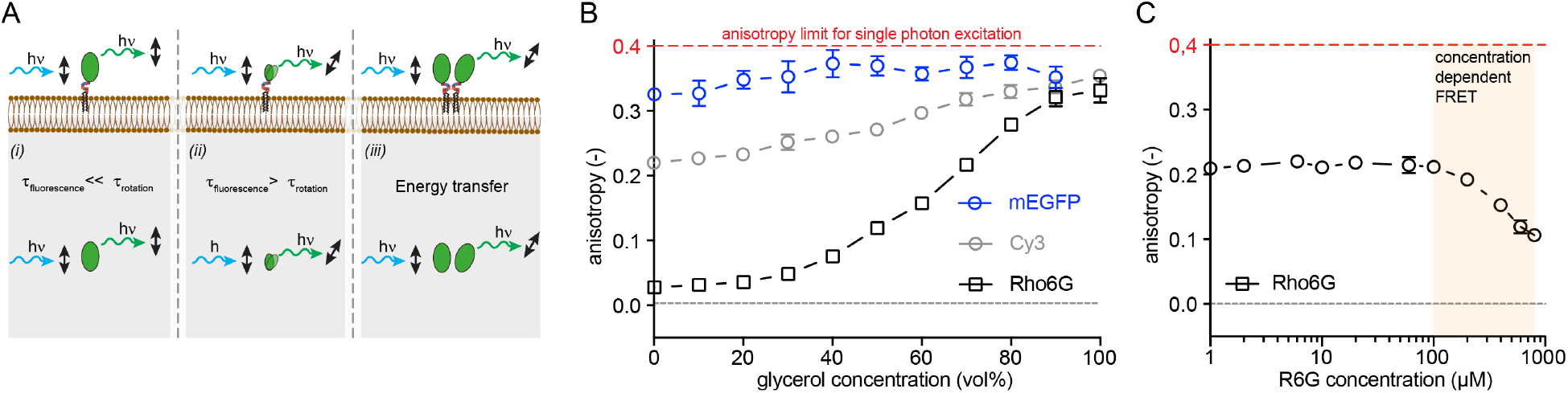
Concept and experiments displaying examples of polarized fluorescence emission anisotropy. **(A)** Fluorophores that have their dipole aligned with the excitation polarization get excited through photoselection. ***(i)*** A fluorophore with limited rotational mobility during its fluorescence lifetime, such as typical fluorescent protein, will emit a photon that has a polarization aligned to the excitation polarization. ***(ii)*** Smaller molecules can rotate much faster and depolarise their emission with respect to the excitation polarization. ***(iii)*** If the energy is transferred to a nearby fluorophore the resulting emitted photon will be depolarised with respect to the excitation polarization, because the second fluorophore is more likely to be differently oriented with respect to the first fluorophore. **(B)** Experimental data showing the effect of fluorescence lifetime and rotational time on the steady state emission anisotropy measurement. Anisotropy increases as the rotational timescale is slowed down though increasing the viscosity of the solution by increasing the glycerol content (x-axis, from 10^−3^ Pa·s to 1.412 Pa·s) or due to a size increase of the fluorophore (EGFP versus Rho6G; Mw of 26.9 kDa versus 0.8 kDa resulting in a *ϕ* of 14 ns versus 0.12 ns). The influence of fluorescence lifetime is exemplified by the higher anisotropy of Cy3 compared to Rho6G due to the shorter lifetime of Cy3 (τ=0.3 ns versus τ=3.43 ns). **(C)** Experimental data displaying the effect of molecular proximity on the anisotropy measurement. Concentration dependent anisotropy of Rho6G in a 70% glycerol solution (∼2.3·10^−2^ Pa·s) contains two regimes: (1) a regime determined only by rotational diffusion and (2) a regime where Rho6G is undergoing increasing collisional homo-FRET upon an increasing concentration (above 100 mM).

For fluorophores that can freely rotate, the extent of depolarization with respect to the excitation is described by the Perrin equation that relates the fluorescence anisotropy to both the fluorescence lifetime and the rotational diffusion time of the fluorophore (Perrin, 1926) (Figure 1Aii):

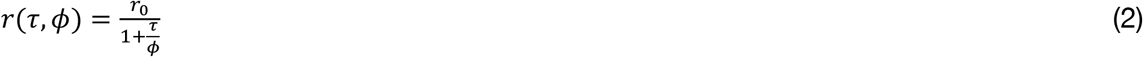

where *τ* is the fluorescence lifetime of the fluorophore and *ϕ* is the rotational time constant. For the small fluorophore Rho6G, its lifetime of 3.43 ns is much larger than its rotational diffusion in water (*ϕ*∼0.12 ns). In essence this means that the emission dipole of an excited Rho6G molecule has had the time to randomly re-orient many times before emitting a photon, resulting in a measured anisotropy close to 0 (Figure 1B). An increased anisotropy is obtained for fluorophores that either have a reduced fluorescence lifetime comparable to its rotational timescale like for example, Cy3, (*τ*∼0.3 ns, Figure 1B) or an increased rotational timescale for example, EGFP, (*ϕ*∼14 ns, Figure 1B).

Increasing the viscosity of the solution increases the rotational diffusion time of a fluorophore. Figure 1B shows that increasing the fraction of glycerol in the solution results in an increase in the anisotropy for all three fluorophores (Rho6G, Cy3 and EGFP). Note that the measured anisotropy of Rho6G almost completely covers the full dynamic range expected from an ensemble of molecules rapidly rotating with respect to its lifetime to being randomly oriented in the static limit where the rotational diffusion is very slow compared to the lifetime of the fluorophore. The emission anisotropy values range from 0 to 0.35, likely limited through the instantaneous depolarisation due to the angular difference discussed above. Although not usually taken into account it is worth to consider the refractive index of the medium. The lifetime of many fluorophores is sensitive to changes in refractive index (Strickler and Berg, 1962; Suhling *et al*., 2002) (*τ* ∝ *η*^−2^; where *η* is the refractive index). Taking a look at the equation 2, it is evident that as refractive index increases, the anisotropy will also increase. Changes in the refractive index may come about by an increase in macromolecular crowding, which is a common state in intracellular conditions (Boersma *et al*., 2015; Berg *et al*., 2017).

The above influences on the value of anisotropy are realized in situations where the fluorophores are dilute and hence far apart from each other. Upon increasing concentration, intermolecular distances reduce, bringing fluorophores within Förster’s radius where Förster’s resonance energy transfer takes place. In this process energy from one fluorophore may be transferred non-radiatively to a nearby fluorophore (Figure 1Aiii). Since the emission dipole of this second fluorophore will be stochastically oriented with respect to the absorption dipole of the first, energy transfer effectively depolarizes the emission from the interacting pair of fluorophores. This depolarization or decrease in anisotropy provides a measure of the efficiency of the energy transfer process:

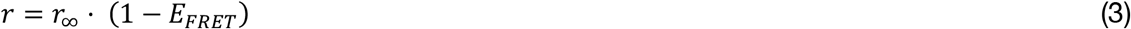

where *r*_∞_ is the anisotropy of the fluorophore at infinite dilution and *E*_*FRET*_ is the energy transfer efficiency. In a dilute solution *r*_∞_ is described by the Perrin equation, and *E*_*FRTE*_ values will rise once the fluorophores in solution start approaching each other at distances, *d*, comparable or closer than the Förster’s radius:

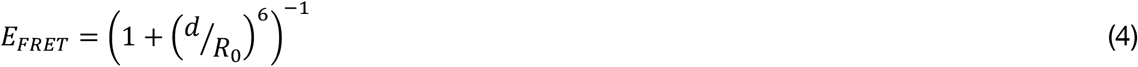

The Förster’s radius, *R*_0_, measured in *Å* is defined as the distance between two fluorophores for which the energy transfer is 50% through (Förster, 1948):

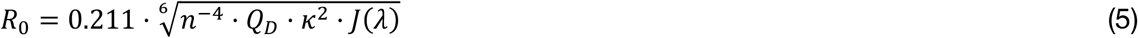

where *J*(*λ*) is the spectral overlap between donor emission and acceptor excitation, *n* is the refractive index of medium in the range of spectral overlap, *Q*_*D*_ is the quantum yield of the donor in the absence of an acceptor and *κ* the orientation factor between the dipole of two fluorophores (normally assumed to be 2/3, but can be measured (Dale *et al*., 1979)).

The effect of collisional homo-FRET can be experimentally realized by measuring the fluorescence emission anisotropy of Rho6G in a 70% glycerol:water solution with increasing Rho6G concentration. Above a concentration of 100 μM the anisotropy of the solution starts decreasing due to energy transfer between fluorophores (Figure 1C), reflecting the fact that at these concentrations a measurable fraction of molecules are within Forster’s radius of each other.

### How microscope polarization characteristics affect the anisotropy measurement

Anisotropy measurements in a microscope are easy to set up and implement on any imaging modality available today, provided a few factors related to preserving the dynamic range of the measurements are fulfilled. Here we outline the factors that relate to this requirement for emission anisotropy measurements conducted in either confocal, TIFR or wide field modalities. In a confocal microscope emission anisotropy measurements have been implemented in point scanning (Bader *et al*., 2009), line scanning (Goswami *et al*., 2008; Ghosh *et al*., 2012), light sheet (Hedde *et al*., 2015; Markwardt *et al*., 2018) or as a spinning disk (Ghosh *et al*., 2012; Gowrishankar *et al*., 2012) system. The wide field microscope has been used in an EPI illumination configuration (Varma and Mayor, 1998; Ghosh *et al*., 2012) or with an appropriate objective in a total-internal reflection (TIRF) configuration (Ghosh *et al*., 2012; Raghupathy *et al*., 2015; Kalappurakkal *et al*., 2019). The heart of the measurement lies in the ability to excite fluorophores with polarized light and collect fluorescence emission with sufficiently high polarization extinction.

To establish the effect of various instrumental parameters we took advantage of the fact that the difference in anisotropy of a 1μM Rho6G solution in water to that measured in 100% glycerol covers almost the entire range of emission anisotropy available for a molecule dissolved in a solvent; from 0, as expected from a molecule rotating much faster than its fluorescence lifetime, to 0.35, where the value of anisotropy is result rotationally averaged distribution of dipoles that are excited by polarized light and have a very low rotational diffusion coefficient. First, the excitation polarization extinction coefficient (*C*_‖_ = *I*_‖_/*I*_⊥_) at the sample plane was gradually reduced from 1500:1 to 1:1 using a λ/4 waveplate (Figure 2A). When the polarization extinction coefficient was reduced below 150:1 the measured dynamic range started decreasing, setting the minimum requirement for the polarization extinction of the excitation path for maximum dynamic range. Placing a high-quality polarizer (e.g. Newport 10LP-VIS-B or MoxTek PFU04C) in a collimated segment of the excitation path before the sample plane will take care that this requirement is met.

**Figure 2.**
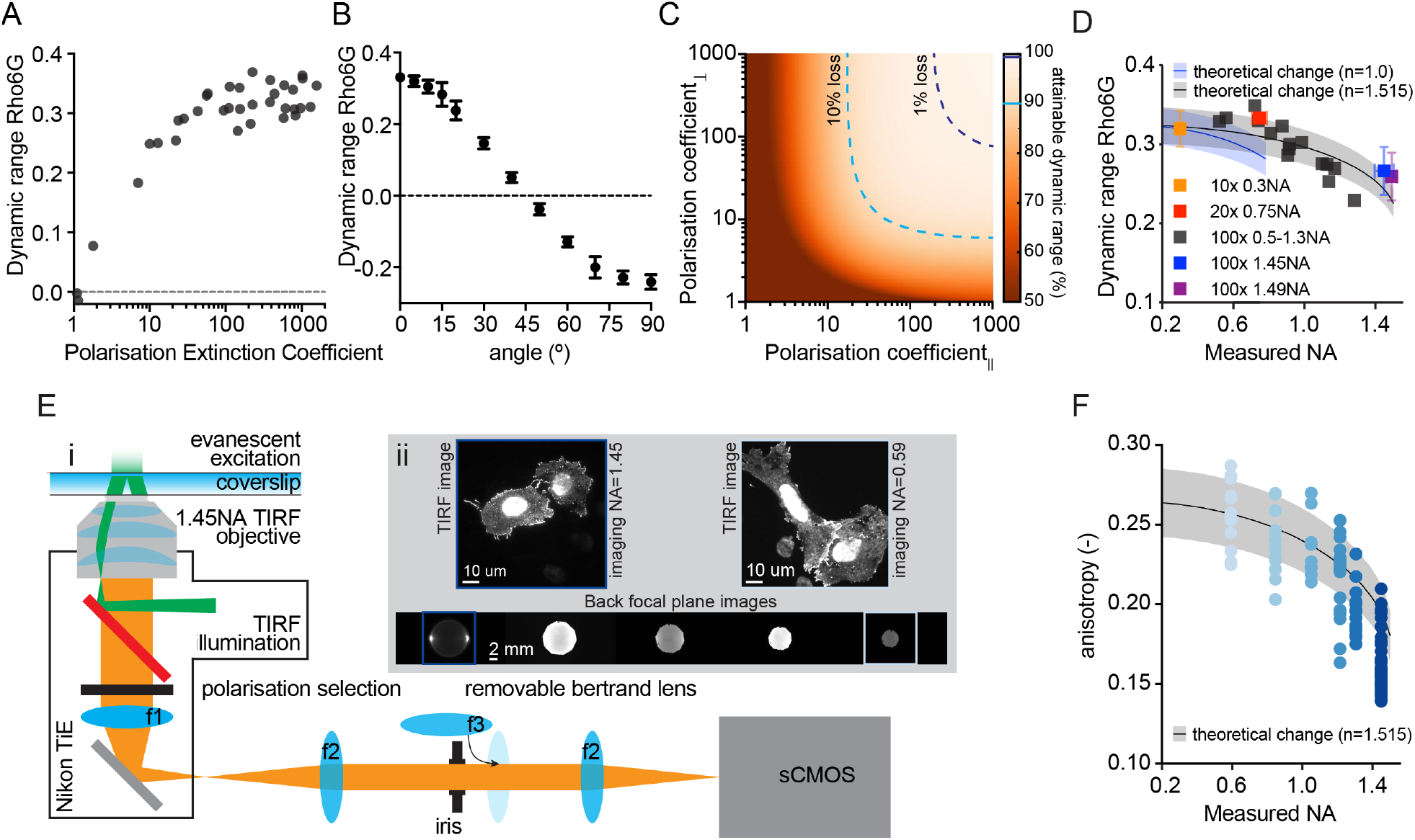
Effects of polarization characteristics on the dynamic range of an anisotropy experiment. **(A)** Lowering the excitation polarization extinction coefficient, by rotating a λ/4 waveplate that is placed in the excitation path, dramatically lowers the dynamic range of an anisotropy experiment. The dynamic range is experimentally determined as the difference between Rho6G dissolved in glycerol versus water. **(B)** The orientation of the polarization of the excitation source can be misaligned with respect to the detection polarization axis by rotating a λ/2 waveplate that is placed in the excitation path. Misalignment significantly reduces the dynamic range eventually flipping the sign of the anisotropy because the parallel and perpendicular detection axes with respect to the excitation polarisation become interchanged above 45º. **(C)** The maximum attainable dynamic range is also determined by the polarization extinction coefficients (or signal-to-noise) for each of the two detector channels. The dynamic range is most sensitive to the extinction coefficient of the parallel channel (x-axis). Nevertheless, to avoid loss of dynamic range (colour scale) both the extinction coefficients should remain ideally above 100:1. The blue dashed lines indicate a 1% and 10% loss of the dynamic range. **(D)** The depolarisation effect associated to high Numerical Aperture (NA) objectives changes the detected anisotropy values and thereby the dynamic range. The NA of a microscope system can be changed by using different NA objectives (different coloured squares) or through changing the aperture size of a variable NA objective (black squares). As expected from theory, the dynamic range decreases with increasing NA. The theoretical change in dynamic range is dependent on the refractive index and was adjusted to the mean (blue and black line for n=1.0 and n=1.515, respectively) and standard deviation (blue and black shaded region) of the measurement using the 10x 0.3NA objective (see Materials and Methods). **(E)** A high NA objective is essential for TIRF illumination and NA reduction should therefore occur at a part of the optical path where excitation and collection are not shared. ***(i)*** Schematic setup used to change the NA selectively at the emission collection side. The focussed sample plane from the microscope tube lens (f1: 200mm) is relayed on a sCMOS camera using two achromatic plano-convex lenses (f2: 100mm). The position and relative opening of the iris in the conjugate back focal plane is monitored using a removable Bertrand lens (f3: 30mm). ***(ii)*** With the Bertrand lens in place the back focal plane can be visualized and controllably constricted using an iris. The resultant change in the detection NA does not affect the evanescent field excitation conditions used to illuminate only the basal cell membrane. **(F)** Steady state anisotropy values of VsV-G-EGFP measured from the basal membrane of cells using TIRF excitation at different collection NAs shows the expected trend of a decreased anisotropy at higher collection NAs. Each point is a single cell. The theoretical change in dynamic range (black line and shaded area as mean and standard deviation, respectively) was adjusted to the mean and standard deviation of the measurement at the lowest NA (see Materials and Methods).

Next, the direction of the maximum excitation polarization (1500:1) was changed with respect to the detection polarizations using a λ/2 waveplate (Figure 2B). A misalignment of less than 5º is desirable for obtaining the maximum dynamic range. Note, negative values of anisotropy arise because the parallel and perpendicular detectors become interchanged after 45º. Because high numerical aperture (NA) objectives scramble the polarization (see below) these polarization calibration measurements were performed using a low NA low magnification air objective (10x, 0.3NA) on an epi-illuminated microscope system.

Finally, on the detection side it is equally important to have a large polarization extinction ratio for each detection channel to obtain the maximum dynamic range in the measurement. The effect of the polarization extinction ratio on each of the detectors can be simulated in terms of bleed through (Figure 2C). For example, a 50:1 extinction coefficient for the perpendicular channel means a 2% contribution from parallel photons. The result is a reduced effective dynamic range of the anisotropy measurement. The most sensitive detector is the parallel channel, and the dynamic range significantly reduces when its extinction coefficient drops below 20:1. One should ideally aim to have an extinction coefficient greater than 100:1 for both channels. These images may be recorded sequentially using two orthogonally oriented high quality polarizers or simultaneously by splitting the emission signal using a polarizing beam splitter and collecting the image on one or two cameras (Ghosh *et al*., 2012). Wire polarizing beam splitters (Moxtek) typically have polarization extinction coefficients on the order of 400:1 on the transmission side and 1:150 on the reflection side with a minimum loss of photons due to absorption. If the extinction coefficient is not satisfactory (i.e. is <100:1) a clean-up polarizer can be placed in front of each detector.

### The influence of the effective numerical aperture on anisotropy measurements

The numerical aperture (NA) of an objective in a microscope system is directly related to the effective angular distribution over which the fluorophores in the sample plane are illuminated. This will also influence the anisotropy fluorescence emission that is collected (in both confocal and EPI/TIRF). The increased illumination and collection angle effectively causes a mixing of polarizations, therefore a lowering of anisotropy. This effect is both dependent on the objective as well as the polarization characteristics of the sample. Even though theoretical correction-factors exist to account for high NA collection (Axelrod, 1979, 1989), reducing the NA of the excitation and emission side will increase the dynamic range of polarization anisotropy measurements (Figure 2D). Reducing the NA of an imaging system may be done by using a lower NA objective or by under-filling a higher NA objective. The former will increase the polarization homogeneity of the excitation field and emission field, while the latter only the excitation field.

Nevertheless, not all microscopy schemes allow for a straightforward reduction in the NA. For objective-based TIRF microscopy, high NA is crucial to obtain the critical angle required for total internal reflection in the excitation path. The axially confined evanescent excitation profile produced by TIRF provides the sensitivity and specificity to image surface localized molecules of interest. It is possible to obtain a pure plane polarised evanescent field by orienting the polarisation of the incoming light perpendicular to the plane of incidence along which total internal reflection occurs (s-polarisation) (Ghosh *et al*., 2012). The mixing of polarization of a TIRF experiment thus mostly occurs during the collection of the fluorescence emission. And in particular at the larger collection angles that are restricted to the outer rim of the objective back focal plane (BFP). These supercritical angle fluorescence photons can be cut off in a conjugate plane of the BFP that is not shared by both the excitation and emission (Figure 2Ei). Physically reducing the collection NA does not perturb the evanescent-confined field of the excitation (Figure 2Eii) and can in fact be used to correct an anisotropy experiment obtained with a high NA objective. We estimated the collection NA by measuring the reduction in the BFP diameter (see Materials and Methods). To understand the effect of reducing the collection NA on an anisotropy measurement we turned to a constitutive trimeric protein complex that expresses at the cell membrane of a cell: EGFP tagged Vesicular stomatitis virus Glycoprotein (VsV-G-EGFP) (Kreis and Lodish, 1986). As expected, the anisotropy of membrane bound VsV-G-EGFP increases upon reducing the collection NA (Figure 2F).

### Error in anisotropy determination

Even in a microscope system that is optimized for the preservation of polarization in the excitation and emission paths, the quantification of fluorescence intensity by a detector has a signal-to-noise that is proportional to the number of photons due to the Poisson statistics of the signal. Using error propagation of the signal collected in the parallel and perpendicular channels allows insight in the relative error directly associated with the anisotropy measurement (Lidke *et al*., 2005):

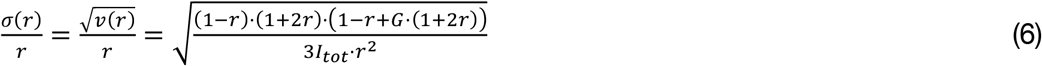

The error is dependent on the anisotropy, *r*, and the total number of detected photons, *I*_*tot*_ Plotting the relative error dependence in pseudocolor on both calculated anisotropy and number of photons (Figure 3A) shows that a relative error below 10% (blue dashed line in Figure 3A) requires a minimum of 400-15000 photons for an anisotropy range of 0.35-0.05, respectively. Using less photons to calculate the anisotropy sharply reduces the accuracy at which the anisotropy can be determined. Practically this means that in a measurement where the number of photons per pixel is limiting, the measurement accuracy of anisotropy will be dependent on the anisotropy value. In such cases the neighbouring pixels or subsequent frames should be summed in order to increase the accuracy of the anisotropy determination at a cost of spatial or temporal resolution, respectively. A Gaussian kernel filter on the two intensity channels also reduces the error in the anisotropy determination (Lidke *et al*., 2005) but a summation is preferred because it will allow a more straightforward dissection of errors simply associated to the number of photons.

**Figure 3.**
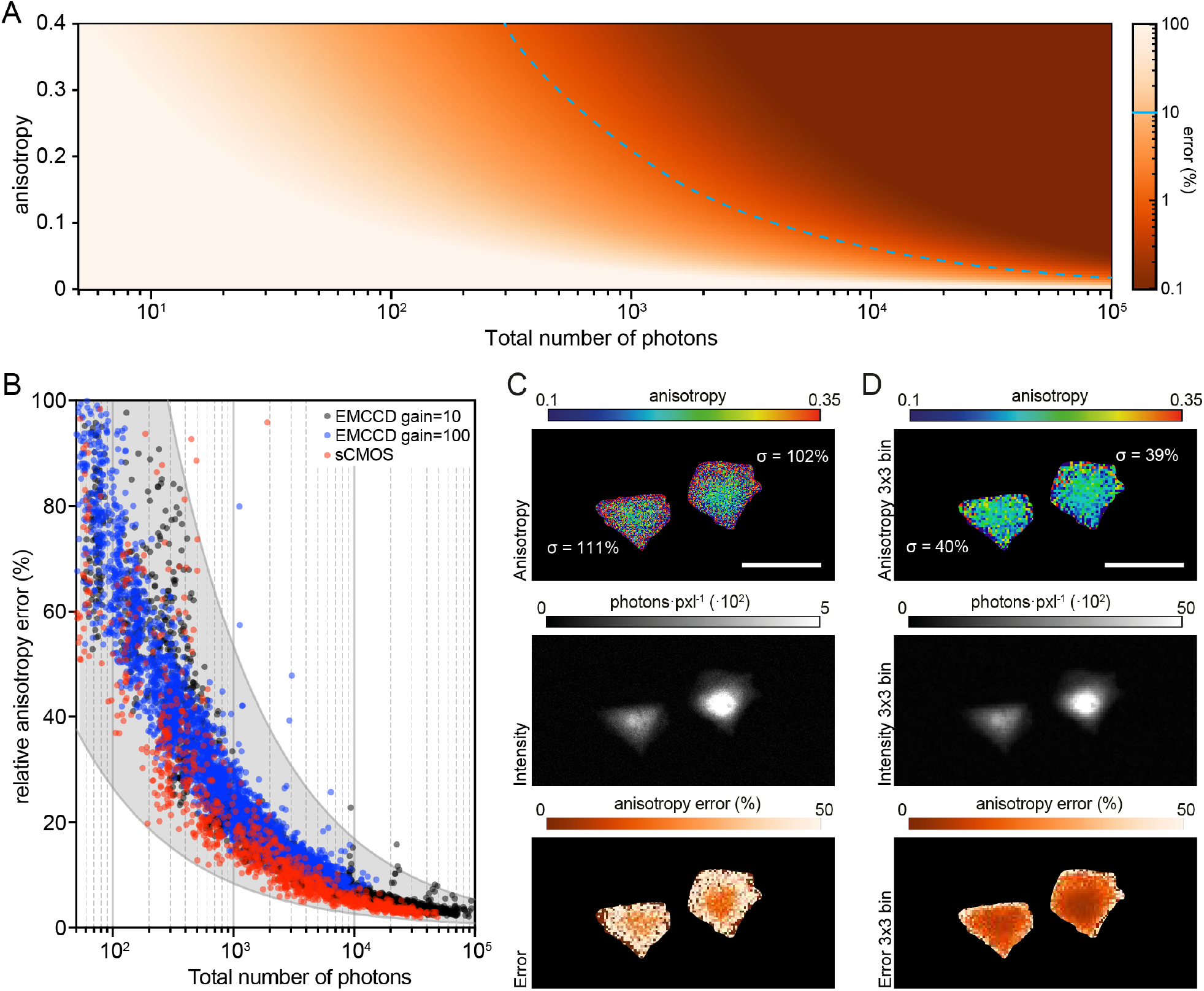
Error in the anisotropy measurement due to photon statistics. **(A)** The relative theoretical error in anisotropy determination (pseudo coloured) is related to both the number of detected photons (x-axis) and well as to the calculated anisotropy (y-axis). The majority of anisotropy measurements from cells collect around 10-1000 photons per pixel constituting a rather large error. This is one of the reasons 10-40 pixel-square regions-of-interest or whole cell regions are collected for quantification. Blue dashed line represents a 10% error. **(B)** The error can also be experimentally measured by recording multiple measurements from an object with a determined anisotropy, such as a 3μm fluorescent bead. The measurement records mean intensity, mean anisotropy and anisotropy standard deviation of each bead. Measured error in the anisotropy from the beads using an sCMOS (red) and an EMCCD with set gain at 100 (blue) or set gain at 10 (black). The errors for all three detectors lie within the grey area associated with the theoretical value. **(C-D)** Images of anisotropy (top panel), total intensity (center panel) and associated pixelwise error (bottom panel) for cells expressing cytosolic EGFP trimers obtained from **(C)** raw and **(D)** 3×3 binned data. Note that increasing the binning of the images in post processing increases the number of photons per pixel and thereby decreases the variability in the anisotropy. Scale bar is 50 μm.

To experimentally determine the error in any anisotropy measurement, the detector must first be calibrated in terms of offset, noise and gain. Depending on the detector there are several methods available to calibrate the system *a prior* (Vliet *et al*., 1998; Huang *et al*., 2013; Lambert and Waters, 2014) and a recent method also allows post-processing of single images (Heintzmann *et al*., 2018). After calibration, a 100-frame anisotropy image series of fluorescent beads was recorded. The intensity and anisotropy trace of a single pixel at the peak position of each bead was used to extract mean and standard deviation (Ferrand *et al*., 2019). The mean intensity, i.e., horizontal axes of the graph (Figure 3B), was increased experimentally by increasing the camera integration time and through binning the intensity images during post-processing (see Materials and Methods). In this way the experimental error can be determined from several detected photons all the way up to 10^5^ photons (Figure 3B). Although the variability in mean anisotropy of the beads is rather high (0.22±0.09), indicating a non-uniform sample, the single bead-associated error remains within the boundaries set by photon statistics for both sCMOS and EMCCD cameras (grayed area in Figure 3B).

### Anisotropy measurements in cells

Expression of proteins from plasmids in cells can encompass a wide range of levels, which for cytosolic proteins would correspond to different cytoplasmic concentrations. Imaging cells that express fluorescent proteins encoded on extra-chromosomal plasmids allow a visual illustration of the variability of anisotropy due to the large range of expression levels possible (pM up to mM; (Milo and Phillips, 2007; Mori *et al*., 2020)). Even a single cell displays the result of the errors due to photon budget on the anisotropy calculations (Figure 3C). The lower number of photons collected from the cell edges (below 100 photons), dramatically increased the variability in the anisotropy calculation. For the pixels within the same cells the relative error in anisotropy is more than 100%. Consequently, binning the number of photons per pixel results in a reduced error in the anisotropy (Figure 3D). This reduction in variability is purely due to the reduced error in the anisotropy calculation associated with using an increased number of photons for the calculation. Therefore, quantification of the signal from high magnification images is typically obtained from selecting sub-cellular regions of interest or the entire cell for lower magnification images (Figure 3D).

To test multiple conditions a common brightness region in the Cell Brightness versus Anisotropy graph must be selected (Figure 4A, in between dashed gray lines). The selection should avoid regions dominated by photon statistic noise, scattering or trivial collisional FRET. Towards the lower brightness end of the curve, noise or scattering will start dominating resulting in a decrease or increase in anisotropy, respectively. The cut-off intensity above which collision FRET becomes the dominating factor for the anisotropy measurement can be identified by a clear change in the slope towards the higher brightness side of the curve (Figure 1C and Figure 4A).

**Figure 4.**
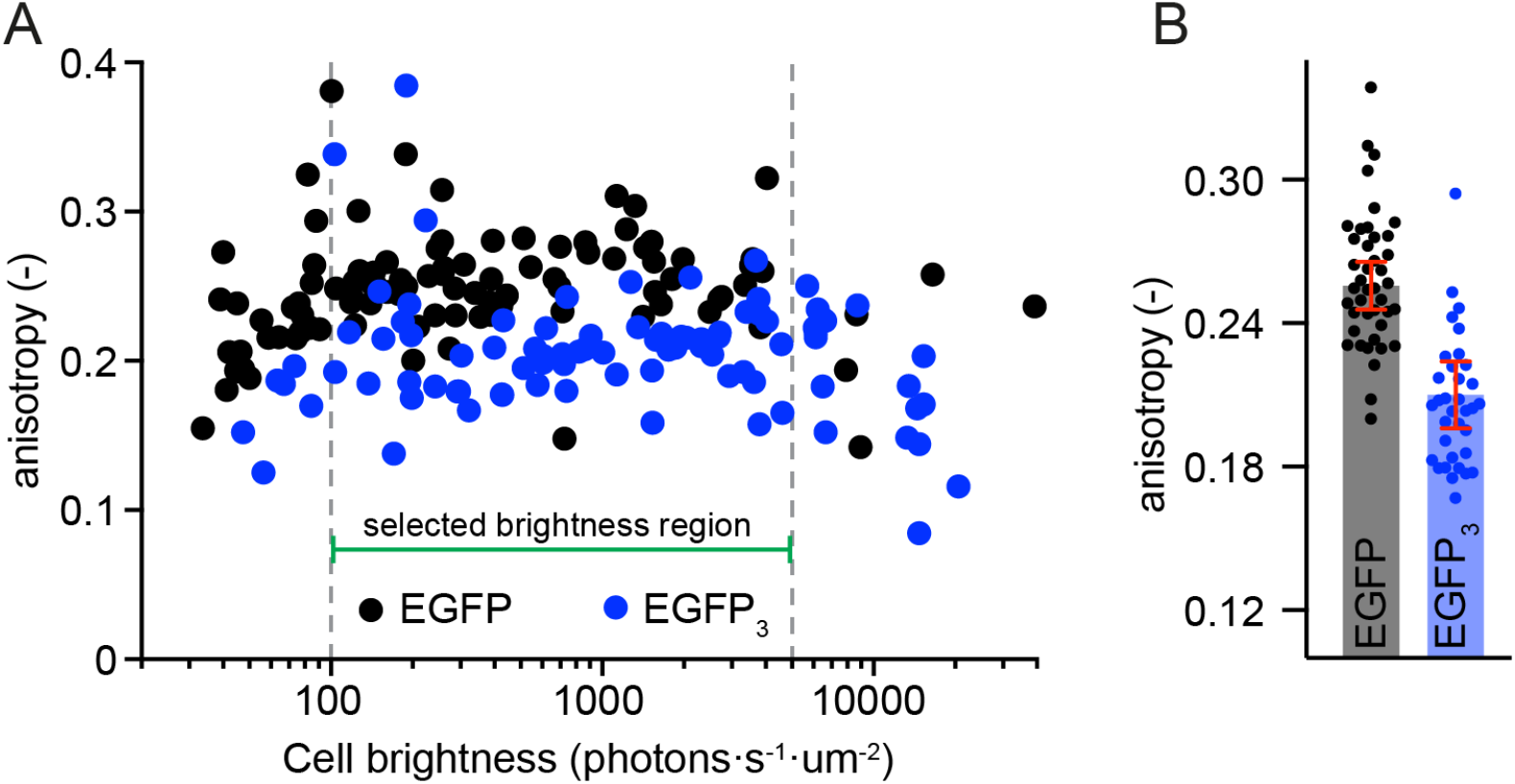
Measurements of GFP anisotropy in living cells. **(A)** Scatter plots of brightness versus anisotropy for cells expressing cytosolic EGFP monomers and cells expressing cytosolic EGFP trimers imaged with a spinning disk confocal microscope. Each point is associated to a single cell. The anisotropy was calculated using the total number of photons collected in the two channels for each whole cell. The brightness was calculated by dividing the total photons collected per cell by its area and the camera integration time. Note that the x-axis will shift depending on the excitation power density and is therefore dependent on the imaging mode (e.g. EPI, TIRF, spinning disk confocal, point-scanning confocal, etc) and excitation power. **(B)** Anisotropy values of cells having expression levels between the indicated levels in panel (A). Please note that in order to compare two or more sets of data the anisotropy values have to be taken from in between two identical limiting values. In this case the boundaries are set by a lower limit from photon statistics and an upper limit after which collisional FRET dominates the anisotropy value.

Collecting the anisotropy values of cells within the selected region shows that the rotational correlation time of EGFP in living cells (17 ns; (Gautier *et al*., 2001; Sharma *et al*., 2004)) results in a measured anisotropy of 0.26±0.03 (Figure 4B). Significant homo-FRET in the trimeric version of cytosolic EGFP gives rise to a lowering of this anisotropy measurement to ∼ 0.21±0.04 (Figure 4B). (It is important to reiterate that the x-axis in Figure 4A is proportional to fluorophore density or concentration). The total number of photons that has been used for the anisotropy calculation is in this case related to both the cell area, its brightness and the camera integration time. The minimum number of photons collected from a single selected cell (3.5·10^4^ photons) within the common brightness region dictates the largest error is ∼ 2-3%, using equation 6. The fact that the standard deviation of the anisotropy values of different cells expressing EGFP (13%) or EGFP-trimer (20%) is larger than the error determined only by photon statistics suggests that there is also a large heterogeneity in the environment of the EGFP and the EGFP-trimer in cells. This may be caused by variation in cell intrinsic properties that change GFP photophysics, or due to the cell-cell differences in the viscosity of the cytoplasm.

### Attenuation of homo-FRET by emitter dilution

While depolarization of emission anisotropy is a sensitive technique to measure molecular interactions and proximity, it is important to make sure the interpretation of the measurement is correct. One way is to always have a proper positive and negative control during the experiment. In the case of cholesterol-sensitive Glycosyl Phosphatidyl Inositol (GPI)- anchored protein nanoclustering, a negative control that disrupted clustering without significantly affecting the rotational correlation times of the labelled GPI-anchored protein was obtained by the removal of cholesterol from the membrane (Sharma *et al*., 2004). However, it is not always known *a priori* for a particular system of interest how the molecules are clustered, rendering the possibility of a negative control in doubt. Especially in situations where it is difficult to clearly distinguish a change in rotational diffusion of the fluorescent probes from a change in homo-FRET, other controls are required. If available, time-resolved anisotropy measurements will be the most direct way to separate these two quantities, since here rotational diffusion times may be deconvolved from the rate of energy transfer in the time resolved anisotropy decay traces (Gautier *et al*., 2001; Sharma *et al*., 2004).

Alternatively, in measurements detailed here, diluting the fluorescent sample while monitoring the steady state emission anisotropy is a relatively straightforward approach to separate these two modes of anisotropy reduction. Photobleaching reduces the number of functional emitters without affecting the rotational dynamics of the fluorophore. Therefore, a steady increase in emission anisotropy with photobleaching the fluorophore is a direct measure of the loss of homo-FRET. Other ways of reducing the number of fluorophores that can participate in the energy transfer process is by the use of quenchers or photoconversion (Ojha *et al*., 2019). It is important to keep in mind that the dark state of fluorophores generated by photobleaching some fluorophores (e.g. GFP in an oxidizing environment, e.g. extracellularly) or even the quenchers themselves should not be able to transfer the energy back to the fluorophores.

To exemplify the effect of photobleaching, cells expressing the cytosolic EGFP trimer were photobleached. These cells displayed a clear increase in the emission anisotropy as the cellular brightness decreased (Figure 5A,C). Cytosolic EGFP monomers also photobleached but in contrast to the trimer there was no change in the emission anisotropy as cellular brightness decreased (Figure 5B,C). Anisotropy was replotted as a function of emitter dilution, which is equivalent to the extent of photobleaching. The measured anisotropy change upon photobleaching per cell shows that while for cells with monomeric EGFP there was a minimal change there was a significant increase in the anisotropy for cells that contained the EGFP trimer (Figure 5D). The measurement for each cell was fitted to a linear curve that provided the anisotropy at full bleaching and the slope. The anisotropy of both the monomeric and trimeric EGFP at infinite dilution were comparable (Figure 5E). In contrast, their initial anisotropies and more importantly the slope, or anisotropy-change upon photobleaching, were markedly different (Figure 5E-F). The absence of a change in anisotropy upon photobleaching (slope = 0.36±0.98·10^−2^) shows that monomeric EGFP did not undergo homo-FRET. On the other hand, for the trimer, a slope of 4.84±1.76·10^−2^ was observed, demonstrating significant and measurable homo-FRET occurring for the EGFP-trimer, as expected. With the reasonable assumption that the anisotropy at point where the trimer and monomer anisotropy values converge represents the anisotropy value at infinite dilution, the FRET efficiency calculated using equation 3, for the EGFP-trimer was 20.1%. This is reasonable considering that for the trimer, the closest distance of approach between two monomers in the trimer is ∼ 3-4 nm (the distance between the fluorophores when GFP is close packed), approximately the same as the Forster’s radius where FRET efficiency is defined as 50%.

**Figure 5.**
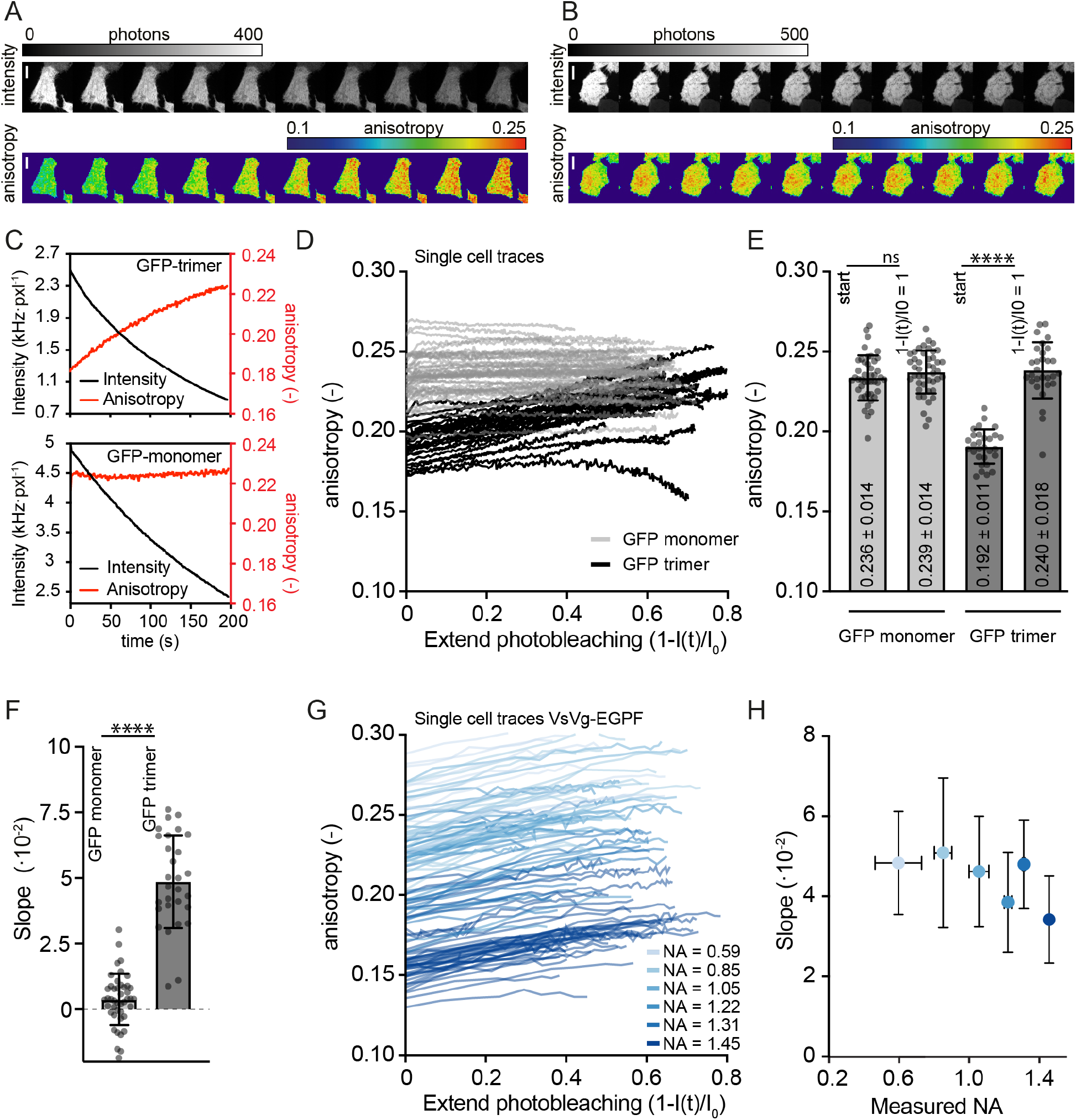
Loss of homo-FRET due to emitter dilution using photobleaching. **(A)** Montage of total intensity (top panel) and associated anisotropy (bottom panel) images of a cell expressing EGFP trimers upon photobleaching from 0 to 200s. Note the increase in anisotropy of the cell as it gradually becomes dimmer upon photobleaching. Scale bar is 10 μm. **(B)** Montage of total intensity (top panel) and associated anisotropy (bottom panel) images of a cell expressing monomeric EGFP upon photobleaching. In contrast to the cell in (A) this cell does not show an anisotropy change even though it is becoming dimmer in time. Scale bar is 10 μm. **(C)** Graphs quantitatively displaying the detected brightness and anisotropy of the cell depicted in panel (A), top, and panel (B), bottom. Note that the brightness values are much higher than the cell brightness associated to the steady state anisotropy measurements of cells in Figure 3C and Figure 4A. This is due to the higher excitation conditions required for photobleaching (4.2 mWatt for photobleaching and 0.35 mWatt for imaging). **(D)** Graphs from various cells relating the extent of photobleaching versus anisotropy for cells expressing cytosolic EGFP monomers (gray lines) and cells expressing cytosolic EGFP trimers (black lines). Each line corresponds to a single cell. The majority of the cells were photobleached to around 20-40% of their original brightness in 100-200 s. Note that the majority of the curves can be represented by straight lines. **(E)** Starting anisotropy and the anisotropy estimated from a straight line extrapolation from the photobleaching curves. The anisotropy from cells expressing the cytosolic EGFP monomer remained largely similar (from 0.236 to 0.239). Cells expressing the EGFP trimer, on the other hand, have a very distinct anisotropy at both ends of the photobleaching (from 0.192 to 0.240). Note that the anisotropy extrapolated to full photobleaching of the EGFP trimer is equivalent to the anisotropy from EGFP monomers. Each point corresponds to a single cell. **(F)** The slopes of the graphs depicted in (D). Each point corresponds to a single cell. **(G)** Graphs from various cells relating the extent of photobleaching versus anisotropy for cells expressing VsV-G-EGFP measured on the basal membrane of cells using TIRF excitation at different collection NAs. Each line corresponds to a single cell. Note that the absolute value of the anisotropy is dramatically altered ranging from 0.13 all the way up to 0.28 upon lowering the collection NA. Nevertheless, all of the examples display a positive slope upon photobleaching reflecting a decrease in homo-FRET of the trimeric complex as it photobleaches. **(H)** The average and standard deviation of the anisotropy versus photobleaching slopes from multiple cells expressing the VsV-G-EGFP trimer at the plasma membrane, measured with a collection NA ranging from 0.59 to 1.45.

Another factor that influences the anisotropy of emission and the dynamic range of its measurement is the NA of the collection optics. To get more insight into how changing the collection NA affects the dynamic range of an anisotropy experiment, the membrane bound trimer VsV-G-EGFP was photobleached (Figure 5G). To reduce the error in anisotropy determination, signal from the plasma membrane of the entire cell was used. Irrespective of the collection NA, emitter dilution corroborates the fact that the low anisotropy measured from VsV-G-EGFP is due to homo-FRET. This reiterates the fact that even though the absolute value of anisotropy is modulated by the optics used in the measurement, the change upon emitter dilution is a rigorous measurement of homo-FRET. Quantifying the slope suggests that there is an additional improvement of the dynamic range upon reducing the collection NA (Figure 5H).

### Red-edge excitation to uncover homo-FRET

The chromophore of most fluorescent proteins undergoes a significant change in dipole moment when driven from the ground state into the excited state. In a situation where the chromophore is surrounded by a polar solvent, solvent movement will accompany the redistribution of the electron cloud during excitation (Lakowicz, 2006). The chromophore will emit a photon from the solvent-assisted relaxed state if the solvent redistributes faster than the excited state lifetime. However, in a fluorescent protein the polar residues that interact with the chromophore have relaxation rates that are much slower compared to the excited state lifetime (Haldar and Chattopadhyay, 2007). This means that a chromophore excited at its main absorption band does not relax to its lowest energetic configuration and will therefore emit a blue-shifted photon. Shifting the excitation towards the red edge of the absorption spectrum will result in photoselection of chromophores that interact more strongly with the surrounding polar residues and are configurationally closer to the final solvent relaxed state (Lakowicz and Keating-Nakamoto, 1984). The emission will consequently also shift more towards the red and is termed red-edge excitation shift (REES) (Demchenko, 2002). REES is additionally associated to a loss of energy transfer in homo-FRET due to the decreased likelihood of resonant coupling between the photo-selected chromophore and its close neighbour. Red-edge excitation can therefore be used to non-destructively probe for the occurrence of homo-FRET for fluorescent proteins (Squire *et al*., 2004).

Imaging cells expressing cytosolic monomeric EGFP displayed no significant difference between anisotropy images calculated from 488 nm and red-edge (514 nm) excitation (Figure 6A). In contrast cells that expressed the EGFP trimer exhibited a significant increase in anisotropy when shifting the excitation from 488 nm to 514 nm (Figure 6B). This is even more striking in the difference image where each cell is predominantly pseudo-colored in red, signifying an anisotropy increase. The cells expressing monomeric EGFP show up in the difference images as pseudo-coloured in both red and blue, indicating that they contain regions of both increased anisotropy that coexist with regions of decreased anisotropy (Figure 6A). This subcellular heterogeneity is most likely due to the temporal shift of about 3s between the 488 nm and 514 nm anisotropy image because it was obtained using a sequential point-scanning scheme and can be improved by using a different simultaneous collection scheme.

**Figure 6.**
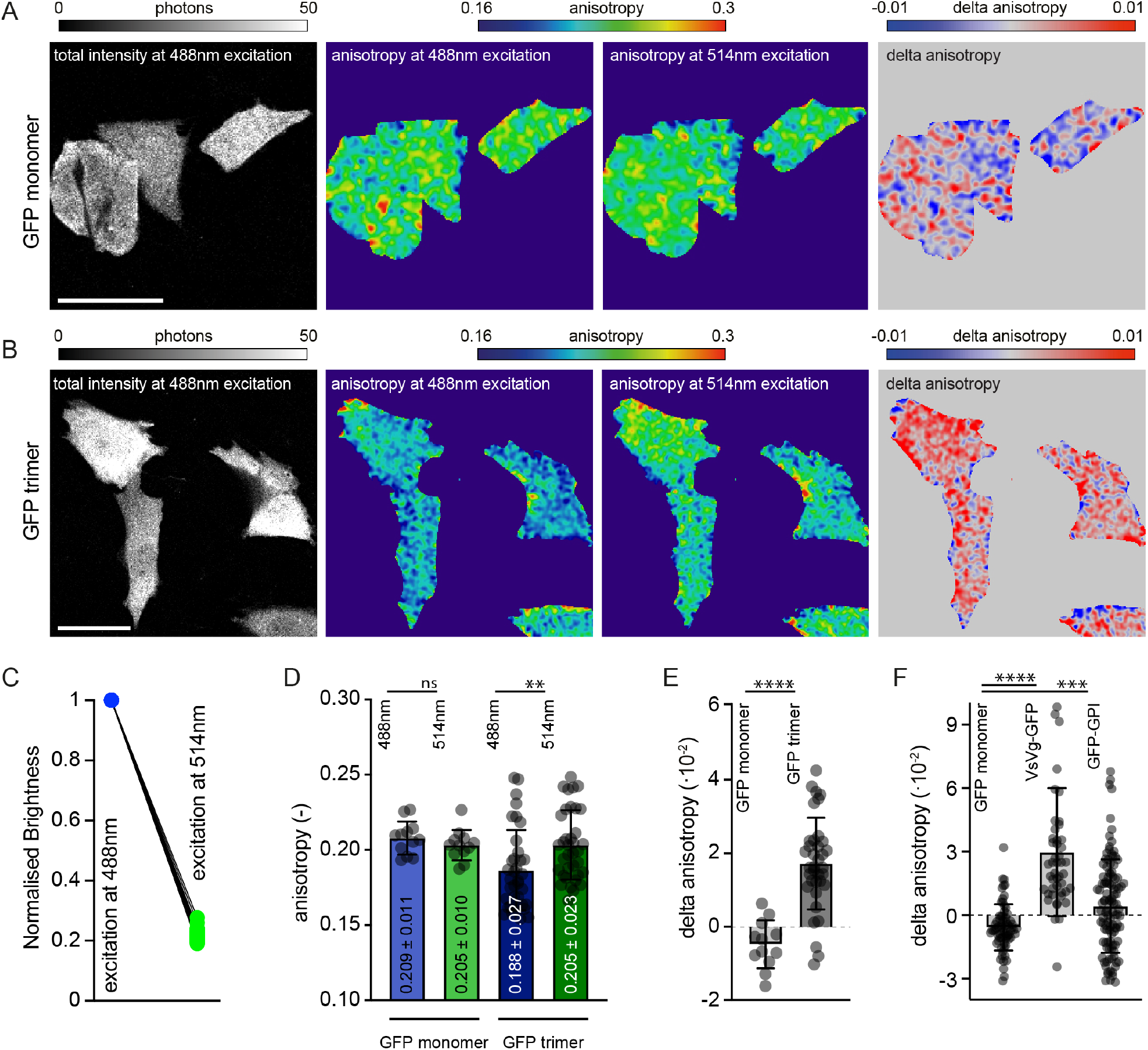
Loss of homo-FRET upon red-edge excitation. **(A-B)** Total intensity, anisotropy at 488nm excitation, anisotropy at 514nm excitation and anisotropy difference upon red-edge excitation of cells expressing **(A)** monomeric EGFP or **(B)** trimeric EGFP. The anisotropy calculations have been obtained from 3×3 binned raw data images. **(C)** Excitation power-dependent decrease in emission intensity upon excitation with 514nm normalized to excitation at 488nm. Since the excitation with 514 nm is at the shoulder of the absorption spectrum of GPF about 5 times more power (or integration) is required to obtain a similar emission intensity, **(D)** Anisotropy measured at 488nm and red-edge excitation of 514nm for cells expressing monomeric or trimeric EGFP. Each point is a single cell. **(E)** The difference in anisotropy between 488nm and 514nm excitation of the same cell expressing either monomeric or trimeric EGFP. Each point is a single cell. **(F)** The change in anisotropy values upon exciting at the red-edge of monomeric EGFP compared to membrane bound trimeric VsV-G-EGFP and EGFP-GPI. Each point is a single cell.

Illumination at 514 nm generates a smaller photon flux per micro watt excitation power as compared to a 488 nm illumination, as expected by the ensemble absorption spectrum of EGFP (Figure 6C). To avoid differences in the error of the anisotropy calculation the laser power was increased when exciting at the red-edge of 514 nm, to obtain similar photon counts. To reduce the error even more the intensity from the entire cell was used to calculate the anisotropy. Red-edge excitation causes a significant increase in the cell-wide anisotropy of cells expressing the EGFP trimer in contrast to the lack of change for cells expressing the EGFP monomer (Figure 6D). Quantifying the change reveals a slightly negative change for monomeric EGFP (−0.47±0.65·10^−2^) versus a positive anisotropy change of 1.71±1.24·10^−2^ for the trimeric EGFP (Figure 6E). Next, red-edge excitation performance was tested on two membrane-bound proteins: the trimeric VsV-G-EGFP and mEGFP-GPI. The GPI-anchored protein is known to have a 20% fraction form small nanoclusters (Sharma *et al*., 2004; Zanten *et al*., 2009). A 3.55±3.02·10^−2^ anisotropy change was measured for VsV-G-EGFP with respect to monomeric EGFP and a 1.01±2.20·10^−2^ change for mEGFP-GPI (Figure 6F). This shows that red-edge anisotropy loss as a measure for homo-FRET is sensitive enough to detect the low amount of clustering of GPI-AP in a sea of randomly diffusing GPI-AP monomers (Sharma *et al*., 2004).

## Discussion

In this manuscript we have outlined methods and caveats associated with the measurement of emission anisotropy derived from the polarized excitation of fluorophores in an imaging mode. We define parameters of optical instrumentation in terms of its influence on the excitation polarization and detection of polarized emission. The role of SNR in terms of number of photons detected also contributes significantly to measurement errors, and ways to mitigate these have been examined. Indeed, taking into account all considerations, the detection of emission anisotropy with sufficient accuracy allows a detection of homo-FRET, permitting a very high-resolution measurement of nanoscale molecular interactions in livings cells.

One of the major challenges associated with homo-FRET detection is that the change in emission anisotropy may have origins in processes other than energy transfer. In fact, a straightforward implementation of steady-state emission polarization anisotropy will not allow an immediate distinction between the effect on energy transfer or rotational changes in the probe. Therefore, the experiments need to be complemented with their proper controls and calibrations. Alternatively, for a time-resolved measurement of the emission polarization, the effects of both energy transfer and rotation can be precisely measured. In fact, the rate of anisotropy decay observed in time-resolved anisotropy measurements due to homo-FRET is equivalent to the rate of energy transfer (Gautier *et al*., 2001), and the ability to deconvolve the fraction of species undergoing this decay informs us about the fraction of species undergoing FRET (Clayton *et al*., 2002). It is worth reiterating that the dynamic range of time-resolved homo-FRET measurements is larger compared to using two distinct fluorophores to measure molecular mixing (Tramier *et al*., 2003). The reason for this is that instead of measuring a small change in the lifetime of the donor fluorophore, time-resolved anisotropy measures the changes due to the appearance of a fast time-scale due to homo-FRET, that is of the order of sub-nanoseconds as compared to rotational effects on anisotropy which are on the order of tens of nanoseconds, especially for fluorescent proteins (Volkmer *et al*., 2000; Sharma *et al*., 2004). However, even while time-resolved measurements are more accurate in determining the extent of energy transfer, the high costs associated with the instrumentation and extensive analysis of the signals may render this approach somewhat less attractive.

Measuring changes in emission polarization anisotropy upon photobleaching, photo-switching or red-edge excitation provide direct access to the homo-FRET population in a steady-state anisotropy set up. Red-edge excitation and the associated anisotropy loss is an attractive and non-destructive alternative to measure clustering. However, this method is crucially dependent on the embedding of a highly polarizable fluorophore in a rigid environment. Therefore, its use in biological contexts will likely be restricted to some fluorescent proteins or labelling strategies that incorporate the chromophores inside a protein pocket (Haldar and Chattopadhyay, 2007) or within a lipid environment (Chattopadhyay, 2003). Another practical consideration is the availability of a high power selective laser source to precisely excite at the red-edge of the fluorophore excitation spectrum. Nevertheless, if all the conditions are met, the method is a powerful tool to probe real-time clustering, as a ratio of anisotropies at these two wavelengths.

The treatment of the effect of NA on polarized detection initially detailed by Axelrod (Axelrod, 1979, 1989), and also documented here (Figure 2F) have largely focused on polarization mixing at large excitation and collection angles. However it should be noted that presence of an interface has additional influence on the emission dipole collection especially at higher NAs (> 1.2) (Oheim *et al*., 2020). This effect is termed super-critical emission collection, and it also decays as a function of distance from the cover slip (Bourg *et al*., 2015). This effect becomes increasingly important for fluorophores of which the emission dipoles are oriented perpendicular to the glass coverslip. The photons emitted by these fluorophores will get equally distributed between the two orthogonally oriented polarization channels and can become detected above super-critical angles at distances less than ∼500nm from the coverslip. Both fast rotation and homo-FRET will permit emission from fluorophores with dipoles oriented perpendicularly to the glass surface and its quantitative influence on anisotropy measurements therefore warrants further exploration.

Finally, the fluorescence imaging modalities used to obtain homo-FRET are still diffraction limited. This means that the measurement of homo-FRET indicates inter-molecular mixing within the respective diffraction limited area, but it does not provide information about the amount of clusters nor their sizes. Careful measurements of the shape of the photobleaching curve, coupled to detailed simulations are likely to give the structure factor and shape of the fluorescent ensembles that undergo FRET, as well as the fraction of molecules undergoing FRET (Sharma *et al*., 2004; Rao and Mayor, 2005; Heckmeier *et al*., 2020). However, even though it is possible to get an estimate of both of these numbers there is no access to the spatial distribution of the fluorophores in the diffraction limited spot. In addition, although the absence of homo-FRET may rule out molecular scale proximity of the probes, it does not exclude the possibility of the association of the probes in a larger complex where individual fluorophore are spaced far enough apart that they do not engage in FRET. Another limitation is the fluorophore size itself which sets the limit on the extent of FRET than may be observed. The sizes of fluorescent proteins (3 nm) prevents the closest approach of fluorophores to approximately this distance, and considering that the Förster’s radius for most fluorescent proteins is around 5 nm (Patterson *et al*., 2000), FRET efficiencies greater than 40% are precluded (Piston and Kremers, 2007).

To gain more insight high resolution homo-FRET imaging can be complemented by indirect techniques based on correlational movement in single particle tracking experiments (Low-Nam *et al*., 2011), number-brightness methods (Digman *et al*., 2008; Cutrale *et al*., 2019) or fluorescence cross-correlation spectroscopy (Bacia *et al*., 2006). More direct measurements of the actual distance between the proteins of interest can be obtained in fixed cells. Spatial patterns can be observed using the super-resolving power of the electron microscope (Prior *et al*., 2003) or super-resolution fluorescence techniques (Hell, 2007).

It is important to note that under conditions when fluorophores are orientationally restricted this will interfere with the measurement of homo-FRET. In an extreme case, if the entire fluorophore population is oriented, even if they are within Forsters’ radius, and are capable of energy transfer, their excitation with polarized excitation will not result in a detectable loss in emission anisotropy. This implicitly means that it would be necessary to ascertain that the ensemble of fluorophores under question does not exhibit an emission anisotropy that is sensitive to the angle subtended with the polarized excitation. For example rhodamine-labelled phalloidin molecules decorating an actin filament, exhibit an emission anisotropy that varies with the angle that the filament subtends with the axis of polarized excitation. Therefore homo-FRET between rhodamine-phalloidin molecules will be poorly detected in case the filament is decorated with a high density of fluorophores. In fact, such a measurement on the same microscopy systems calibrated for sensitive emission anisotropy measurements can be used to measure the orientation of the fluorophores, provided the fluorophores are restricted in their movement and align themselves with respect to a structure or protein of interest. Using such a method, the orientation of actin filaments (Cruz *et al*., 2016; Rimoli *et al*., 2022), integrin receptors (Nordenfelt *et al*., 2017) and nuclear pore complexes (Kampmann *et al*., 2011) within a cell have been visualized.

In conclusion, the large dynamic range and sensitivity provided by emission polarization based FRET microscopy makes it a technique that is exquisitely suitable to measure small changes in nanoscale clustering of proteins. Here we have provided a practical guideline that will allow the identification of errors associated with the measurement and outlined several of the caveats that need to be considered whilst making such measurements. In addition, we have indicated several ways to ascertain that the measured value corresponds to the property of interest, e.g., nanoclustering.

## Acknowledgements

T.S.v.Z. acknowledges an EMBO fellowship (ALTF 1519-2013) and the NCBS Campus fellowship. S.M. acknowledges a JC Bose Fellowship from the Department of Science and Technology (Government of India) and support from a Welcome Trust/Department of Biotechnology, Alliance Margdarshi Fellowship (IA/M/15/1/502018).

## MATERIALS AND METHODS

### Fluorophores and beads

Rhodamine 6G (Sigma) and Cy3 (GE healthcare) were dissolved in milliQ water at 1mM and further diluted with milliQ or glycerol (Merck) to obtain the indicated concentrations of fluorophores at the required water:glycerol content. EGFP was dissolved in PBS was further diluted in PBS and glycerol keeping the EGFP concentration at 100nM. A coverslip with 3 μm beads (BD biosciences) was prepared by depositing 50 μl of a 10^−4^ dilution followed by drying.

### Cells

CHO-K1 cells were cultured in phenol-red free HF12 (Himedia) supplemented with 10% FBS (Gibco) and 1% antibiotics (Gibco). Cells were plated on uncoated glass dishes 48 hours before and transfected with EGFP-N1, EGFP-EGFP-EGFP-N1 or VsV-G-EGFP 12-16 hours before the experiment using Fugene6 (Promega). CHO cells stably expressing GFP-GPI were obtained earlier (Sharma *et al*., 2004). To clear the golgi and ER content GFP-GPI and VsV-G-EGFP expressing cells were exposed to 50 μg/ml of cycloheximide (Sigma) for a total of 3-4 hours before the experiment.

### Anisotropy measurements, calibrations and error determination

Fluorophores or beads were excited using either a laser (Agilent MLC100, 488nm line) send in via the TIRF arm or a LED (CoolLED pE-300-ultra with a Chroma 480/20x excitation filter) via the EPI arm of a Nikon TiE microscope. Both excitation sources had a polarizer (Moxtek PFU04C) at the collimated region of their beam path, were directed to the objective via a dichroic (Semrock Di03 R405/488/562/635) and had polarization extinction coefficients of 1500:1 (laser) and 365:1 (LED). Emission light was collected using the same objective and filtered for emission (Semrock 520/35) and polarization (Newport 10LP-VIS-B) right after the dichroic but before the tubelens of the microscope. The emission light was subsequently sent via a Cairns optical splitter to the camera. The camera was either an EMCCD (Photometrics, Evolve delta) or an sCMOS (Photometrics, Prime95B) and the polarization images were taken sequentially. Both cameras had been calibrated for noise and pixel gain values before the experiments using documented methods (Vliet *et al*., 1998; Huang *et al*., 2013; Lambert and Waters, 2014).

Calibration measurements were obtained using a 10x 0.3NA objective that was focused 5-15 um inside the solution of a 10ul droplet on the coverslip. To change the extinction coefficient of the excitation beam a λ/4 wave plate was placed right after the polarizer in the path carrying laser light. Turning the λ/4 waveplate changes the polarization extinction coefficient, which was measured before each measurement. In a similar fashion the orientation of the excitation placing and rotating a λ/2 waveplate right after the polarizer in the excitation path altered polarization. sCMOS camera integration times were 200-8000 ms and a laser excitation power of 1.5-2 mWatt ensured between 500-1200 photons per pixel. The error was further reduced by using the average of 5-10 recorded frames, each containing >10^5^ photons per region of interest.

Once converted into photo-electrons the images were aligned with respect to each other, either using the image itself or the alignment matrix from separately imaged subdiffraction beads. Any postprocessing on the raw intensity such as binning or smoothing was performed at this stage. Subsequently, the total number of photons from each polarization in different regions of interest were extracted and together with the measured G-factor (0.99±0.02) used for anisotropy calculation using equation (1). The regions of interest (ROIs) were; a 100×100 pixel area at the center of field of view for solution images, the peak position of a bead, or an entire cell. In some cases also the mean number of photons was used to determine the brightness.

Bead images under LED illumination with a 10x 0.3NA objective were used to experimentally determine the errors associated with the anisotropy calculation. Single pixel peak positions of beads were obtained through a 2-D Gaussian fitting on maxima found in the average intensity image of a 100-frame time-series. This allowed the extraction of a temporal trace from both polarizations, which was used to calculate the anisotropy using the G-factor (1.06±0.02) and equation (1). The mean and standard deviation of the anisotropy over the 100-frame trace was then used in combination with the average total number of photons per frame to generate a single point in Figure 3B. To cover a large region within photon-budget space both camera integration time (10-1000ms) and post-process pixel binning (0-5 pixels) was used for all camera settings. LED excitation was used because the non-coherent nature avoids interference issues that might significantly affect frame-to-frame intensities of single beads and the power was kept at 0.1 mWatt resulting in negligible photobleaching.

### Changing Numerical Aperture

The numerical aperture of the EPI/TIRF system was changed in two ways. The first was by using different objectives: 10x 0.3NA air, 20x 0.75NA air, 100x 0.5-1.3 variable NA oil, 100x 1.45NA oil, and a 100x 1.49NA oil objective. The collar of the variable NA oil objective changes the filling of the back focal plane by opening and closing an internal iris and thereby alters the effective NA of the objective. The G-factor slightly changed depending on the objective and was corrected in each experiment. The second manner of changing the NA was done in a conjugate image of the back focal plane that was not shared by excitation and emission. For this the emission path was relayed outside the microscope in a 4-f configuration using two 100mm achromatic plano-convex lenses (Melles Griot). Assisted by a removable Bertrand lens (30 mm bi-convex, Thorlabs) an iris was placed and aligned at the conjugate back focal plane of a 100x 1.45NA TIRF objective. Adjusting the diameter of the iris, *d*_*iris*_, changes the NA of the detection path:

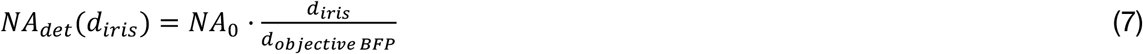

where *NA*_0_ is the NA of the objective, *d*_*objective BFP*_ is the diameter of the objective back focal plane. Anisotropy images were obtained and analyzed as before. The G-Factor (0.99±0.02) remained unaltered upon changing the collection NA.

### Photobleaching in TIRF

In the same set up as described above cells expressing the trimeric VsV-G-EGFP at the plasma membrane were photobleached in TIRF through continuous exposure to 10 mWatt 488nm laser power in TIRF. Camera integration times were 100-300ms and whole cell basal membrane ROIs were taken, excluding large highly intense spots. Photobleaching rates were measured to be 49±16 s^-1^ and independent of detection NA settings.

### Photobleaching in Confocal

Cells expressing cytosolic EGFP-monomer or EGFP-trimer were placed on a CSU-W1 spinning disk confocal. After collimation the excitation beam was sent through a polarization filter (Moxtek PFU04C), a dichroic (Semrock T405/488/568/647) and via the 50 μm pinhole Nipkow spinning disk (4000 rpm) to the objective. Emission was collected with the same objective and from the dichroic sent to two EMCCD detectors (Andor Life888) via a polarizing beam splitter (Moxtek FBF04C) and an individual filter (Chroma ET525/50m) for each camera. The polarization extinction coefficient for 488nm excitation was 3600:1 at the back focal plane and the G-Factor 1.30±0.07. With a 100x 1.4NA objective and 4.2 mWatt power at the back focal plane the photobleaching rate of EGFP in cells was 196±50 s^-1^. Both polarization images were obtained simultaneously at 1 Hz with 50-150 ms integration time and both cameras set at a measured EM gain of 140. To ensure continuous photobleaching the laser was kept on during the photobleaching and typically reached a photobleaching fraction of 0.6-0.8 after 150-200 s. Whole cell ROIs were used and cells containing saturated pixels were not taken further for analysis. Steady state anisotropy single cell analysis of cells expressing cytosolic EGFP-monomer or EGFP-trimer (Figure 3) were also measured on the CSU-W1 spinning disk confocal in a similar fashion as described above, with the exception of using lower laser power (0.35 mWatt) and longer camera exposure times (150-500 ms).

### Red-Edge anisotropy

Cells expressing the indicated construct were imaged on a Zeiss LSM780 with a 40x 1.2NA water objective using highly polarized (>1500:1) 488nm and 514nm excitation. Excitation and emission were separated using MBS488 for 488nm excitation or MBS458/514 for 514nm excitation and the polarization was sequentially selected using orthogonally oriented polarizers. After identical band pass filter settings (518-562nm) the photon stream was detected using the 32-array GaAsP detector that was set at pseudo photon counting mode. To ensure similar detection conditions the power of the 514nm excitation was increased to 162 μWatt as compared to 37 μWatt power used for 488nm excitation and the pinhole together with the objective collar position were optimized before the experiment. GFP excitation under these conditions is still within the linear regime. The pixel size was set at 415nm, the pixel dwell time at 6.3 μs and photobleaching of the sample was minimal. Due to the point scanning mode there was a temporal difference between polarizations of 2.7 s and a wait time between two excitation conditions of 5.2 s. Switching the sequence of acquisition had no influence on the cell measurements. The G-Factor of the system was determined using a 100nM FITC solution and was 1.154 for 488nm excitation and 1.157 for 514nm excitation. Because the FITC chromophore is freely rotating in a solution with fast solvent dynamics the polarization of the emission will explore all orientations irrespective of the excitation conditions.

### Theoretical calculations

In order to theoretically estimate the effect of signal-to-noise at the detectors on the anisotropy measurement a contamination was introduced in the calculation of the anisotropy:

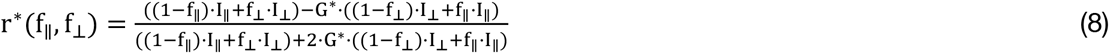

with f_‖_ the fraction of I_‖_ leaking into the perpendicular channel, f_⊥_ and I_⊥_ the reverse. f_‖_ and f_⊥_ are the inverse of their polarization extinction coefficients. This calculation has the underlying assumption that no photons are lost during the detection. The G-factor is also affected following:

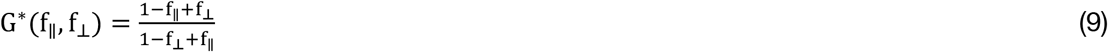

The dynamic range using the two experimental extremes for fluorophores with aligned excitation and emission dipoles, which are r = 0 (I_‖_ = I_⊥_) and r = 0.4 (I_‖_ = 3I_⊥_) can now be calculated with respect to the polarization contamination.

Next, the theoretical error of the anisotropy measurement had been described by Lidke et al. (Lidke *et al*., 2005) and was used to calculate the relative anisotropy error dependence on both the anisotropy and the total number of photons used, see equation 6.

Finally, the effect of the numerical aperture (NA) of the system on the anisotropy measurement is calculated following earlier documented equations (Axelrod, 1979, 1989; Piston and Rizzo, 2008). These relate how a high NA lens collects fluorescence emitted into three-dimensional space, (x, y, z), where the z-direction (I_*z*_) of the sample coordinate plane is defined as the parallel (*I*_‖_) plane of the detection. Detection of dipole projections from the other directions than results in:

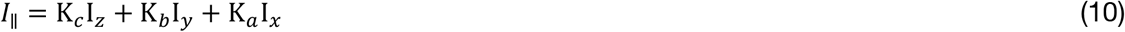

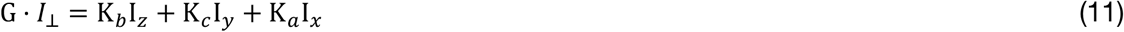

where the normalized weighing factors, K_*a,b,c*_, are defined following Axelrod (Axelrod, 1989):

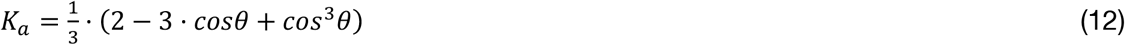

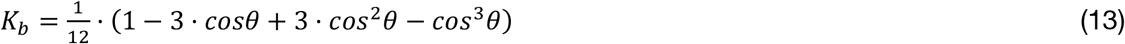

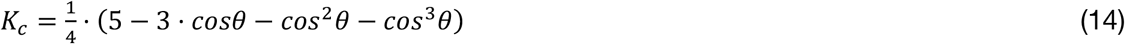

the angle, *θ*, comes from the numerical aperture, NA, via:

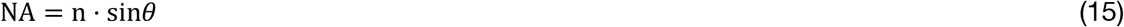

where *n* is the refractive index. Note that for isotropic samples I_*y*_ = I_*x*_ and that for very small angles (low NA) the weighing factor K_*C*_ approaches 1 while K_*a,b*_ are close to 0. In this situation the dipole projections in sample space (*z, y*) follow the detection planes (‖,⊥) For the calculations of the theoretical graphs in Figure 7A,C the anisotropy values measured with the lowest NA (mean ± *σ*) were used as a starting point. Note that Figure 7A has both air (*n* = 1) and oil (*n* = 1.512) objectives.

